# Beating the gold standard: A review of *Mycobacterium tuberculosis* lysis using bead beating and the need for standardization

**DOI:** 10.1101/2025.07.07.663234

**Authors:** Jason D Limberis, John Z Metcalfe

## Abstract

Bead beating is widely used for mechanical lysis of *Mycobacterium tuberculosis*, a bacterium with a highly resistant, lipid-rich cell wall. Despite its status as a de facto gold standard for mycobacterial lysis, there is no standardized protocol for bead beating, resulting in significant variability across studies. We conducted a literature review of 73 studies, identifying 38 with explicit mycobacterial bead beating protocols. Our analysis revealed heterogeneity in bead types, sizes, device models, and operational parameters, with 37% of studies failing to report critical details such as lysis speed. We experimentally assessed the impact of key variables—tube type, bead quantity, and device settings—on lysis efficiency using qPCR of *M. tuberculosis* DNA. Results showed that even minor changes, such as tube shape or bead volume, can significantly affect DNA yield. These findings underscore the need for standardized bead-beating protocols to improve reproducibility and comparability. Future efforts should prioritize developing consensus methods tailored to sample type and analytical application.

## Main Text

Researchers commonly use bead beating to mechanically disrupt cells by agitating small beads at high speed. It is considered the gold standard Mycobacterial cell lysing due to their robust, mycolic acid-rich cell walls that are resistant to chemical disruption. Recently, the Gates Foundation Global Grand Challenges stated in their Enhancing HIV and TB Diagnosis application call: “The solution must break open the MTB cells (>50% lysis efficiency compared to mechanical lysis via bead beating or sonication) without damaging target DNA.”^1^ But there is no standardization for bead beating with different bead sizes, tube sizes and types, bead beating devices, shaking speed (measured in RPM, Hz, oscillations.min^−1^or m.s^−1^), duration, rest time between cycles, and total number of cycles.

We performed a literature review on PubMed on June 29th, 2025, using the search term “(bead-beater) OR (bead-beating) OR (beadbeater) OR (bead beater) OR (bead beating) OR (bead homogenizer) OR ribolysis OR (MP bio) OR (MP Biomedicals) OR MPBio OR beadbug OR (BioSpec Products) OR (Bullet Blender) OR (Mechanical lysis) OR (bead ruptor) AND (mycobacterium OR tuberculosis OR m tuberculosis OR Mtb)”. This search yielded 73 papers, of which 38 included mycobacterial bead beating protocols (**Table 1**).

**Table 1:** Bead beating parameters for *Mycobacterium tuberculosis* lysis reported in the literature. NA indicates not applicable (single cycle protocols); “Not stated” means the parameter was not specified in the original study.

| Bead material | Bead size (mm) | Bead volume | Tube type | Bead beater | Speed<sup>⁹</sup> | Time (s) | Rest (s) | Cycles | Sample volume (μl) | Ref |
| --- | --- | --- | --- | --- | --- | --- | --- | --- | --- | --- |
| Zirconium | 0.1 | Undisclosed | 1.5ml screwcap polypropylene tubes (Sarstedt) | Mini-beadbeater (model not stated; Biospec Products) | Not stated | Not stated | Not stated | Not stated | Not stated | <sup>5</sup> |
| Zirconium | 0.1 | 300mg | Not stated | Mini-beadbeater (model not stated; Biospec Products) | Not stated | 30 s or 60, 120, 180, or 300 | NA | 1 | Not stated | <sup>6</sup> |
| Zirconium | 0.1 | Not stated | Not stated | Beadbeater (Biospec Products) | Not stated | 120 | Not stated | 3 | 350ml | <sup>7</sup> |
| Zirconium | 0.1 | Prefilled lysing matrix B tubes | 2ml screw cap | Mini-beadbeater-1 (Biospec Products) | 4800 RPM (max speed) | 30-45 | 90 | 10-12 | 500-750 | <sup>8</sup> |
| Zirconium | 0.1 | Prefilled lysing matrix B tubes | 2ml screw cap | Not stated | Not stated | 45, 30 | 300 | 1,1 | 1000 | <sup>9</sup> |
| Zirconium | 0.1 | 200μl | Not stated | TissueLyzer II (Qiagen) | 30Hz (max speed) | 180 | NA | 1 | 200 | <sup>10</sup> |
| Zirconium | 0.1 | 300μl | 2ml screw cap | TissueLyzer (Qiagen) | 30Hz (max speed) | 100 | Not stated | 2 | 600 | <sup>11</sup> |
| Zirconium | 0.1 | 75mg | Not stated | Mini-beadbeater (model not stated; Biospec Products) | Not stated | 60 | 60 | 3 | 300 | <sup>12</sup> |
| Zirconium | 0.1 | Not stated | Not stated | Not stated | 30Hz (max speed) | 30 | 60 | 5 | 1000 | <sup>13</sup> |
| Zirconium | 0.1 | 100μl | Not stated | Mini-beadbeater (model not stated; Biospec Products) | Not stated | 30-480 | NA | 1 | 500 | <sup>14</sup> |
| Zirconium | 0.1 | Not stated | Screw cap microfuge tube | Mini-beadbeater (model not stated; Biospec Products) | Maximum speed | 10 | 60 | 5 | Not stated | <sup>15</sup> |
| Zirconium | 0.1 | 200μL | Not stated | Fastprep-24 Classic (MP Bio) | 4.5m.s<sup>-1</sup> | 60 | 60 | 3 | 1000 | <sup>16</sup> |
| Zirconium | 0.1 | 250mg | 2ml screw cap | Mini-beadbeater-1 (Biospec Products) | 4800 RPM (max speed) | 60 | 30 | 3 | 625 | <sup>17</sup> |
| Zirconium | 0.1 | 400μl | Not stated | Mini-beadbeater (model Not stated; Biospec Products) | Not stated | 60 | 60 | 10 | Not stated | <sup>18</sup> |
| Zirconium | 0.1 | 200μl | Not stated | Beadbeater (model Not stated; Biospec Products) | Not stated | 60 | NA | 1 | 800 | <sup>19</sup> |
| Glass | 0.1 | 100μl | Not stated | Mini-beadbeater-16 (Biospec Products) | 3450 oscillations.min<sup>-1</sup> (max speed) | 120 | NA | 1 | 500 | <sup>20</sup> |
| Glass | 0.1 | 800mg | 2ml screw cap | Mini-beadbeater (model not stated; Biospec Products) | Not stated | 60 | 60 | 3 | 800 | <sup>21</sup> |
| Glass | 1 and 0.1 | Half fill the Eppendorf tube<sup>⁹</sup> | Not stated | Mini-beadbeater-16 (Biospec Products) | High<sup>⁹</sup> | 300 | NA | 1 | 200 | <sup>22</sup> |
| Not stated | 0.1 | 600mg | Not stated | Bead beating module (Alere Technologies) | 2400 RPM (unknown max speed) | 240 | NA | 1 | 600 | <sup>23</sup> |
| Glass | 0.15 | 250mg | Not stated | Mini-beadbeater-16 (Biospec Products) | 3450 oscillations.min<sup>-1</sup> (max speed) | 60 | Not stated | 2 | Not stated | <sup>24</sup> |
| Zirconium | 0.5 | 100μl | Not stated | Mini-beadbeater (model not stated; Biospec Products) | Not stated | Not stated | Not stated | Not stated | 200 | <sup>25</sup> |
| Zirconium | 0.5 | Not stated | Not stated | Model Not stated (Biospec Products) | 50 RPM | 90 | NA | 1 | Not stated | <sup>26</sup> |
| Zirconium | 0.5 | Not stated | Not stated | Digital disruptor genie (Scientific Industries) | 3000 RPM (max speed) | 180 | NA | 1 | 50 | <sup>27</sup> |
| Zirconium | 1.4 | Prefilled lysing matrix D tubes | 2ml screw cap | FastPrep-24 Classic (MP Bio) | 4.0m.s<sup>-1</sup> | 60 | NA | 1 | 200 | <sup>28</sup> |
| Zirconium | 0.5 | 300mg | 2ml screw cap | MagNA Lyser instrument (Roche Diagnostics) | 5000 RPM | 30 | Not stated | 3 | Not stated | <sup>29</sup> |
| Zirconium | 0.5 | 150mg | 2ml screw cap | Vortex-Genie 2 (Scientific Industries) | 2700 RPM (max speed) | 60 | NA | 1 | Not stated |  |
| Glass | 0.5 | 200μl | 2ml screw cap | Mini-beadbeater 16 (Biospec Products) | 2500 oscillations.min<sup>-18</sup> | 300 | NA | 1 | 500 | <sup>30</sup> |
| Silica | 0.5 | Not stated | Not stated | Mini-beadbeater 8 (Biospec Products) | Not stated | 120 | NA | 1 | Not stated | <sup>31</sup> |
| Zirconium | 1.4 | Prefilled lysing matrix D tubes | 2ml screw cap | Not stated | 6.5m.s<sup>-1</sup> (unknown max speed) | 60 | Not stated | 2 | 400 | <sup>32</sup> |
| Not stated | Not stated | Not stated | Screw cap Eppendorf tube | Qbiogene bead beater (Qbiogene*) | 6.5m.s<sup>-1</sup> (unknown max speed) | 45 | NA | 1 | Not stated | <sup>33</sup> |
| Not stated | Not stated | Not stated | Not stated | FastPrep FP120 (Qbiogene*) | 5.5m.s<sup>-1</sup> (unknown max speed) | 30 | NA | 1 | Not stated | <sup>34</sup> |
| Not stated | Not stated | Not stated | Not stated | FastPrep 120 (Savant Instruments) | 6.5m.s<sup>-1</sup> (unknown max speed) | 45 | NA | 1 | Not stated | <sup>35</sup> |
| Not stated | Not stated | Not stated | Not stated | TissueLyzer II (Qiagen) | 30Hz (max speed) | 600 | NA | 1 | 575 | <sup>36</sup> |
| Not stated | Not stated | Prefilled lysing matrix B | 2ml screw cap | Fastprep-24 Classic (MP Bio) | 6.5m.s<sup>-1</sup> (max speed) | 60 | NA | 1 | 1000 | <sup>37</sup> |
| Zirconium | Not stated | Not stated | Not stated | Vortex (Not stated) | Not stated | 900 | NA | 1 | 1000 | <sup>38</sup> |
| Not stated | Not stated | 300mg | Not stated | Fastprep-24 Classic (MP Bio) | 6.5m.s<sup>-1</sup> (max speed) | 60 | Not stated | 2 | 600 | <sup>39</sup> |
| Not stated | Not stated | Not stated | Not stated | FastPrep-24 5G (MP Bio) | 6.0m.s<sup>-1</sup> | 40 | NA | 1 | 708 | <sup>40</sup> |
| Glass | 0.5 | 200μl | 1.5ml Eppendorf tube | FastPrep-2 (MP Bio) | 4000 cycles.min<sup>-1</sup> | 5 | NA | 1 | 700 | <sup>41</sup> |

\*Qbiogene was a subsidiary of MP Biomedicals. <sup>⁹</sup>If the device is set to its maximum speed, this is indicated in brackets. <sup>⁹</sup>This device has a single speed, but this is unknown to most readers. <sup>⁹</sup>Reported as 2500 oscillations.min<sup>-1</sup>; however, the listed model only has a single speed of 3450 oscillations.min<sup>-1</sup>. <sup>⁹</sup>ratio of 1 mm to 0.1 mm beads was 3:2 (v:v).

Our literature review revealed significant heterogeneity in bead beating parameters. Zirconium predominated the bead material (77%), while 0.1mm beads were the most common bead size (63%). Researchers most frequently used Mini-beadbeater devices from Biospec Products (60%), followed by the MP Bio FastPrep series (20%). Treatment durations typically ranged from 30-300 seconds, and sample volumes ranged from 50-1000μl (excluding one study with a sample volume of 350ml). We found critical gaps in parameter reporting. Thirty-seven percent of studies failed to report speed parameters, and those that did used inconsistent units, making direct comparisons difficult. This lack of standardization hinders reproducibility and cross-study comparisons.

Multiple factors that we found to be poorly reported likely affect bacterial lysis efficiency, including tube shape, tube orientation, bead volume, liquid volume, bacterial count, bead type and size, and oscillation speed (**Table 1**). To test this hypothesis, we selected representative parameters for which we had device access. We used *Mycobacterium tuberculosis* MC^2^ 7901 at 500 000 CFU/ml in 10mM Tris-HCl buffer and performed bead beating using 0.1mm zirconium/silica beads (Biospec Products, USA) as outlined in **Table 2**, with six replicates per condition. We centrifuged the tubes at 16 000RCF for 5 minutes following bead beating, and 2µl of supernatant was used for a qPCR assay in quadruplicates. We performed reactions in 10μl volumes using Luna Universal qPCR master mix (NEB, USA) targeting *atpE* with the following primers and probe: “GTAACGCGCTTATCTCCGGT”,“AGTATGCCGCCTCAACCAAA”,and“5’-VIC-CAACCCGAGGCGCAAGGGC-MGB-3’”.We calculated relative quantities using FastPrep-24 at 6.5m.s?^1^ for two cycles of 45s in a 1.5ml tube containing 200mg 0.1mm Zirconium beads and 600µl of sample as the reference condition. The relative quantification formula was: RQ = 2^(−ΔΔCt), where ΔΔCt = ΔCt(test samples) − ΔCt(reference samples) and ΔCt = Ct(target gene) − Ct(reference gene). We calculated differences between groups using pairwise Wilcoxon tests with Bonferroni correction for multiple comparisons.

**Table 2:**
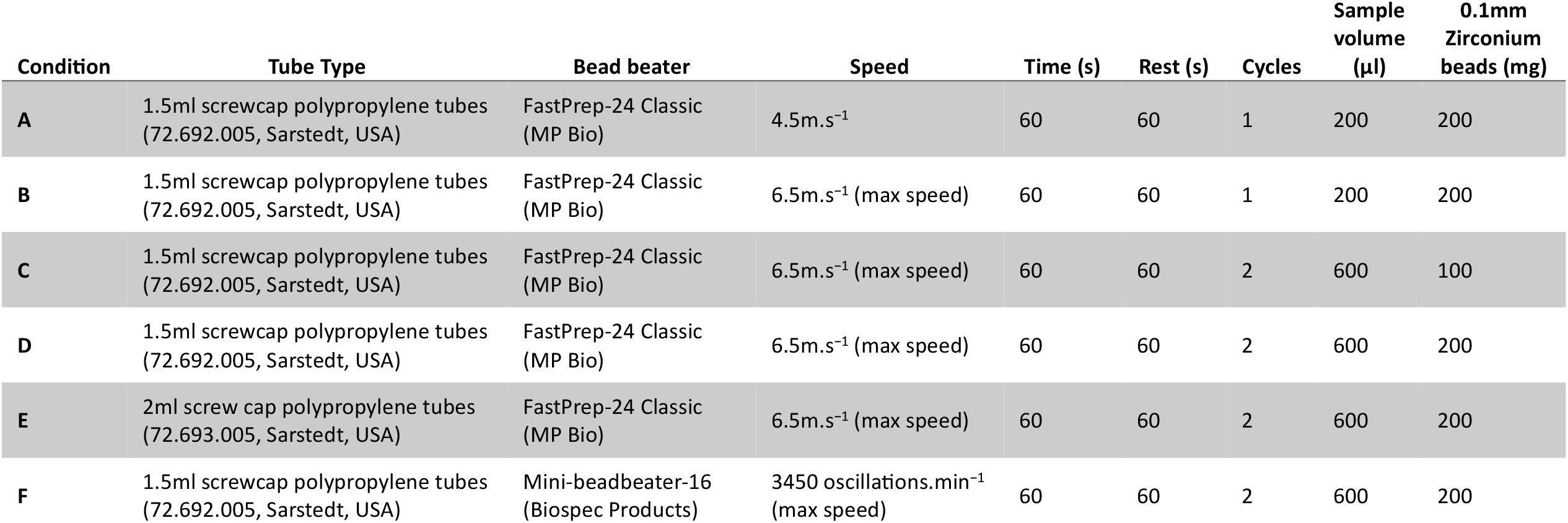
Bead beating conditions used in this study.

Several samples showed substantially different DNA amounts (**Figure 1**). The 2ml tubes yielded a relative quantity of only 0.2 (IQR 0.08, 0.37), probably due to their differing shape compared to the reference tubes. Somewhat counterintuitively, bead beating with fewer beads increased the detectable DNA nearly 4-fold. This finding highlights how seemingly minor parameter variations can dramatically affect results. Our findings demonstrate that many factors affect bead-beating efficiency. Furthermore, this technique lacks standardization beyond mycobacterial applications, and extraction methods introduce significant bias in microbiome profiling. Zymogen developed the ZymoBIOMICS Microbial Community Standard to address standardization challenges.^2^ This mixed microbial community contains well-defined ratios of three easy-to-lyse Gram-negative bacteria (*Pseudomonas aeruginosa, Escherichia coli*, and *Salmonella enterica*), five tough-to-lyse Gram-positive bacteria (*Listeria monocytogenes, Bacillus subtilis, Lactobacillus fermentum, Enterococcus faecalis*, and *Staphylococcus aureus*), and two tough-to-lyse yeasts (*Saccharomyces cerevisiae* and *Cryptococcus neoformans*). Zymogen recommends using their 2ml BashingBead tubes containing 0.6ml dry volume of mixed 0.5mm and 0.1mm ZR BashingBead lysis matrix. Their protocol involves bead beating for 1 minute at 6.5m.s^-1^ for five cycles with 5-minute rests between cycles using the MP Bio Fastprep-24 (Classic or 5G) or the Biospec Mini-BeadBeater-16, and for 5 minutes at maximum speed for four cycles with 5-minute rests between cycles using the Biospec Mini-BeadBeater-96 with 2ml tubes. In the product manuals, BioSpec recommends using 0.1mm beads (quantity not reported) and using up to 400mg of wet weight biomaterial per ml of buffer and running for 2-3 minutes using the Mini-beadbeater-16 (single speed of 3450 oscillations.min^−1^); while MPBio recommends using Lysing Matrix A (Garnet matrix and one 6.35mm ceramic sphere), B (0.1mm silica spheres), F (1.6mm aluminum oxide and silicon carbide particles), or J (2mm yellow zirconia beads and 1.6mm aluminum oxide particles) for bacterial lysis and two rounds of 6m.s^-1^ for 45s with a 5min rest period for mycobacterial cell lysis using the FastPrep-24 5G.^3,4^

**Figure 1:**
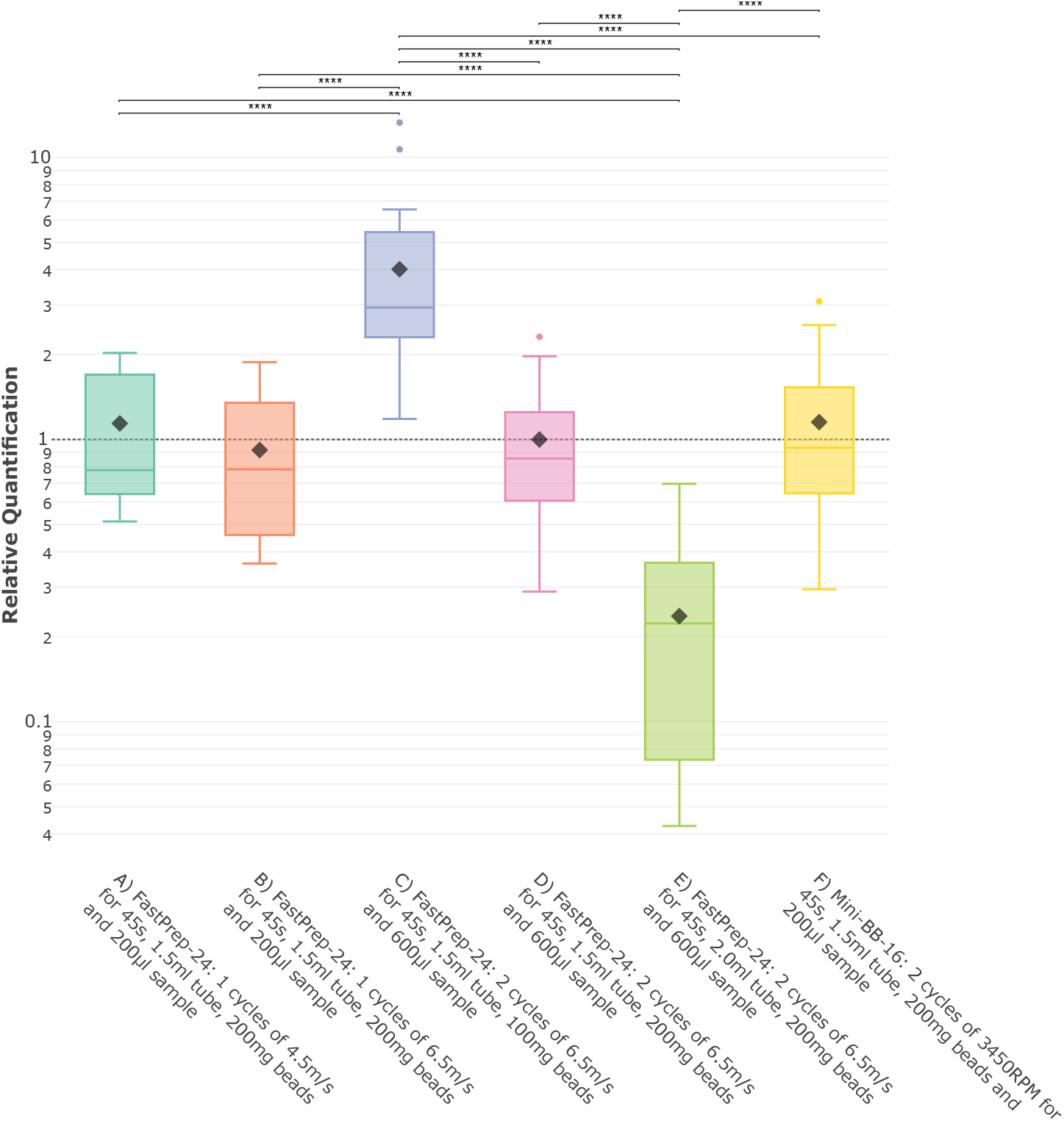
Many variables affect the lysis of *Mycobacterium tuberculosis* by bead-beating. qPCR on lysates from six different bead-beating conditions are shown. We set FastPrep-24 at 6.5m.s^-1^ for two cycles of 45s in a 1.5ml tube containing 200mg 0.1mm zirconium beads and 600µl of sample as the reference condition for relative quantification calculations. Box plots show the distribution of relative quantification values, with the center line representing the median, boxes showing the interquartile range (IQR), and whiskers extending to 1.5 times the IQR and diamonds at the mean. The dashed horizontal line at y=1 represents the mean quantification level of the standard protocol. Statistical significance was determined using pairwise Wilcoxon tests with Bonferroni correction for multiple comparisons (* p ≤ 0.05, ** p ≤ 0.01, *** p ≤ 0.001, **** p ≤ 0.0001).

There is a need to compare and standardize bead-beating protocols for mycobacteria across different sample types (sputum, tissue, blood, environmental samples) and analytical purposes (PCR, sequencing, proteomics). Our findings demonstrate that seemingly minor parameter variations—such as tube geometry, bead quantity, and device selection—can dramatically alter lysis efficiency. Without standardized protocols, researchers cannot reliably compare results across studies or laboratories, hindering progress. Future work should thus focus on developing consensus protocols that account for the multiple variables affecting lysis efficiency while maintaining reproducibility across different laboratory settings and analytical applications.

